# Effects of the Coenzyme Q_10_ Analog 6-Bromo-ubiquinone (6-Br-Q_0_C_10_) on Mammalian Cell Growth

**DOI:** 10.64898/2026.02.28.708723

**Authors:** Brandon Yu, Claire Yu, Peiran Lu, Daniel Lin, Xuejuan Tan, Yong Cheng, Kunhong Xiao, Chang-An Yu

## Abstract

Synthetic 6-Br-Q_0_C_10_ has been shown to exhibit a partial electron transfer activity of native coenzyme Q in the isolated mitochondria. It reduces energy coupling efficiency by approximately 30%, suggesting that it may be useful in modulating cell growth in tissue culture. Whether or not it behaves in the same way in the whole cells, or animal, however, has not yet been fully examined. Recently we have investigated the effect of 6-Br-Q_0_C_10_ across multiple cell lines using three detection methods. Treatment with 6-Br-Q_0_C_10_ reduces cell proliferation in all cell lines tested, with different effectiveness. Obesity-related cell lines were the most susceptible, and a pronounced inhibitory effect was also observed in cancer cell lines. These results strengthen the idea of using 6-Br-Q_0_C_10_ to manage obesity or to retard the growth of rate cancer cells and thus prolonging life.

## Introduction

More than 90% of the energy required to sustain life, growth and the structure integrity of living organisms is generated from a process named oxidative phosphorylation (2-4). This process takes place in the mitochondrial inner membrane of eukaryotic cells or in the cytoplasmic membrane of prokaryotic organisms, via a coupling reaction of two multi-subunit membrane protein complexes: the electron transfer chain complexes (Complexes I (5, 6), II (7, 8), III (9, 10), and IV (11, 12)) and ATP synthase complex (complex V) (11, 12). The electron transfer chain complexes catalyze the oxidation of NADH and succinate generated from tricarboxylic acid (TCA) cycle during the catabolic oxidation of nutrients. This process generates a proton gradient and membrane potential across the inner mitochondrial membrane, which drive the synthesis of ATP from ADP and inorganic phosphate through the action of the ATP synthase complex [Fig. 1].

**Figure 1.**
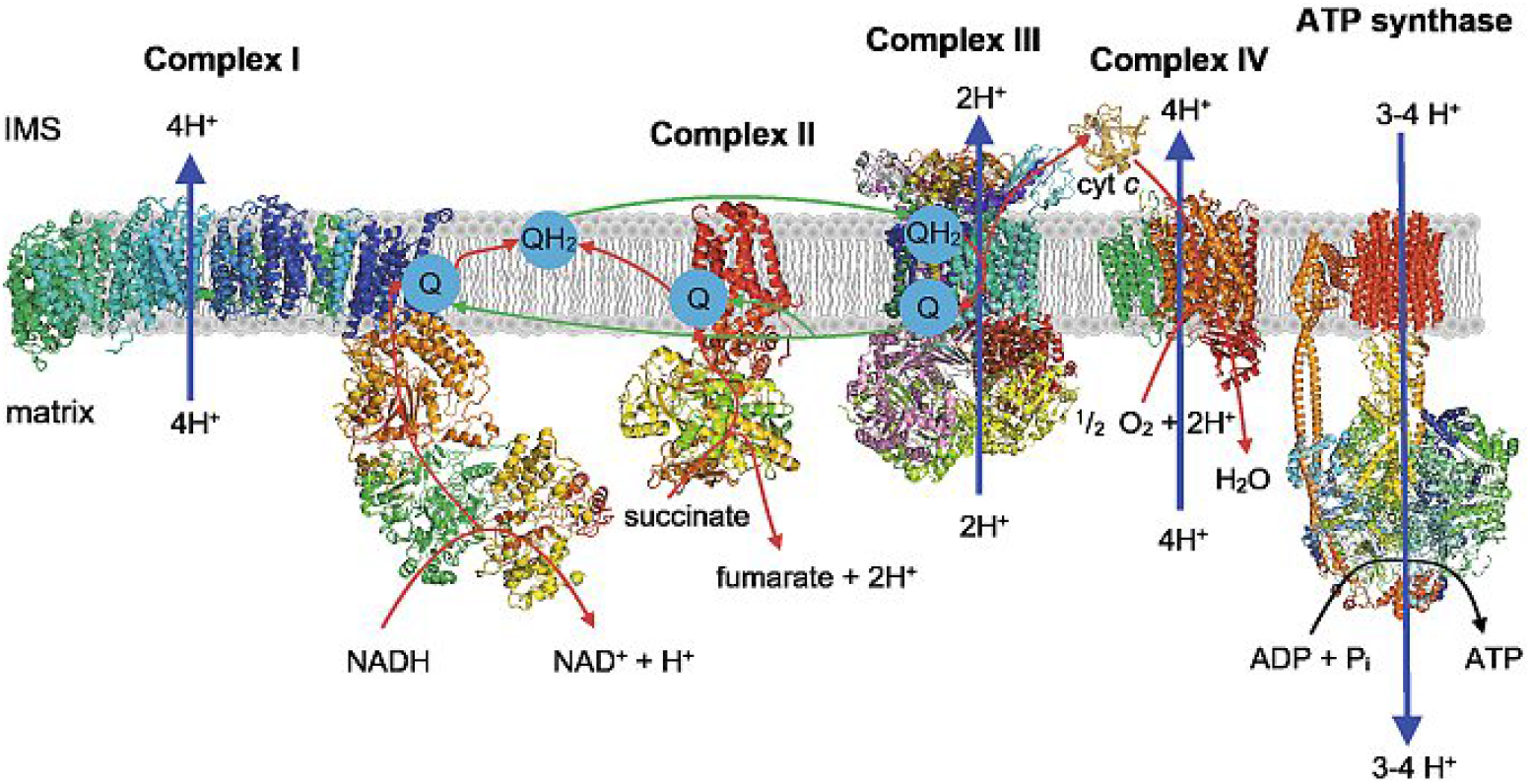
**Electron transport complexes I-IV and ATP synthase (complex V) in the mitochondrial inner membrane** (adopted from *Chimia International Journal for Chemistry* 72, 291-296, 2018) (1).

Among the four of electron transfer complexes, complexes I, III and IV contribute to the generation of the proton gradient, whereas complex II does not directly pump protons across the inner mitochondrial membrane. Coenzyme Q_10_ (2,3-dimethoxy-5-(isoprenyl)_10_-6-methyl-1,4-benzoquinone) (13) (Fig. 2A) mediates electron transfer between Complexes I or II and III. As a key redox component in the electron transfer system, the cellular concentration of coenzyme Q_10_ can be influenced by dietary intake. However, biochemical study of coenzyme Q_10_ is technically challenging due to its low solubility. To address this limitation, researchers in1970s synthesized various coenzyme Q_10_ derivatives with shorter alkyl side chains to increase their solubility while maintaining their electron transfer activity (14, 15). The electron transfer activity of these synthetic derivatives is dependent on the carbon chain length of the alkyl group of coenzyme Q, with the maximal activity reached with a length of 10 carbons, which is much shorter than 50 carbons found in natural coenzyme Q_10_. The most widely used derivative is known as decyl-benzoquinone (DBQ, or Q_0_C_10_) (14), which has a ten-carbon alkyl sidechain. Q_0_C_10_ has been used extensively in the mitochondrial bioenergetic research. Fig. 2B below shows the chemical structure of Q_0_C_10_.

**Figure 2.**
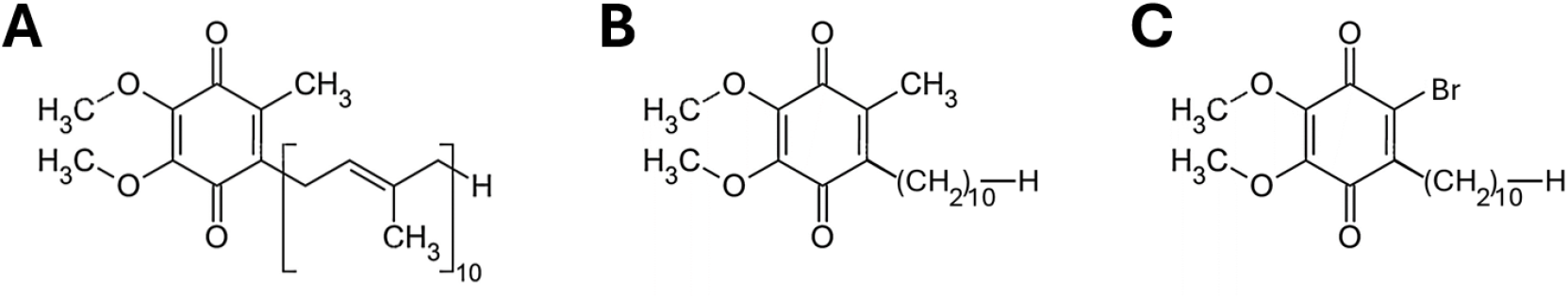
The chemical structure of coenzyme Q_10_ and two derivatives. (A) Q_10_ (2,3-dimethoxy-5-(isoprenyl)10-6-methyl-1,4-benzoquinone), (B) Q_0_C_10_ (2,3-dimethoxy-5-decyl-6-methyl-1,4-benzoquinone), and (C) 6-Br-Q_0_C_10_ (2,3-dimethoxy-5-decyl-6-Br-1,4 benzoquinone).

The spectral and redox properties of Q_0_C_10_ are identical to those of native coenzyme Q_10_ with an absorption maximum at 277 nm and a midpoint redox potential of 110 mV. By replacing the 6-methyl group of the benzoquinone ring of Q_0_C_10_ with a bromine atom, our laboratory had synthesized 6-Br-decyl-benzoquinone (6-Br-Q_0_C_10_) (16). Fig. 2C shows the chemical structure of 6-Br-Q_0_C_10_. Introducing an electron withdrawing group such as bromine should have an effect on the redox potential and UV absorption spectrum. Indeed, it has a higher redox potential than that of Coenzyme Q_10_, by about 30 mV and UV spectral shift by 24 nm. Isolated mitochondria treated with 6-Br-Q_0_C_10_ showed a roughly 30% drop in energy conservation efficiency. This decrease may be due to the loss of energy conservation site II at Complex III in the 6-Br-Q_0_C_10_ treated mitochondria. Similar effects are also observed on the other Halo-CoQ such as 6-Cl-Q_0_C_10_ treated mitochondria.

Whether this reduction in energy coupling efficiency also occurs in intact cells or whole organisms has not yet been determined. Taking advantage of the availability of 6-Br-Q_0_C_10_, we have recently investigated the effect of this compound on cell growth using commercially available cell lines. As expected, 6-Br-Q_0_C_10_ significantly inhibited cell growth. These findings suggest that 6-Br-Q_0_C_10_ might hold therapeutic potential in mitigating obesity or slowing cancer progression, as they are heavily dependent on cellular energy supply.

## Materials and Methods

### Chemical synthesis

Crude 6-Br-Q_0_C_10_ was synthesized as we published previously (16) and available in the laboratory. It was further purified to homogeneity using thin-layer chromatography. The purity of 6-Br-Q_0_C_10_ was verified by its absorption peak at 303 nm, and the concentration was measured spectrophotometrically using a millimolar extinction coefficient of 20 (14).

### Mammalian cell lines

Various mammalian cell lines were used in this study. Information on the cell lines and their corresponding growth medium was listed in **Table 1**. Dulbecco’s Modified Eagle Medium (DMEM) and RPMI 1640 media were purchased from Thermo Fisher Scientific and Sigma-Aldrich, respectively. Bisbenzimide was purchased from Fisher Scientific. CCK-8 reagents were obtained from Boster Bio, and MTT assay kit (ab211091) was purchased from Abcam.

**Table 1:** Characteristics of mammalian cell lines used in this study.

| Cell Line | Cell Type | Origin | Tissues | Culture Medium |
| --- | --- | --- | --- | --- |
| **rMc-1** | Retinal Müller cell | *Rattus* (rat) | Eye; Choroid; Retina | DMEM + 10% FBS |
| **ARPE-19** | Retinal pigment epithelial cell | *Homo sapiens* (human) | Eye; Retina; Retinal pigment epithelium | DMEM + 10% FBS |
| **HepG2** | Hepatocyte | *Homo sapiens* (human) | Liver | DMEM + 10% FBS |
| **L929** | Fibroblasts | *Mus musculus* (mouse) | Subcutaneous connective tissue; Adipose; Areolar | DMEM + 10% FBS |
| **RAW 264.7** | Macrophage | *Mus musculus* (mouse) | Ascites | DMEM + 10% FBS |
| **THP-1** | Monocytes | *Homo sapiens* (human) | Peripheral blood | RPMI 1640 + 10% FBS |
| **HEK293** | Kidney epithelial cell | *Homo sapiens* (human) | Kidney; Embryo | DMEM + 10% FBS |
| **HeLa** | Cervical epithelial cell | *Homo sapiens* (human) | Uterus; Cervix | DMEM + 10% FBS |
| **INS-1** | Insulinoma cells | *Rattus* (rat) | Pancreas; Islet of Langerhans | RPMI 1640 + 10% FBS |
| **ADSC** | Adipose-derived mesenchymal stem cells | *Homo sapiens* (human) | Adipose tissue | MesenPRO RS Medium |

### Cell counting assay

rMc-1, ARPE-19, HepG2, RAW 264.7 and L929 cells were cultured in DMEM with 4.5 g/L glucose supplemented with 10% (vol/vol) heat-inactivated FBS (SH30910.03; Hyclone), and 1x penicillin and streptomycin at 37°C and 5% CO_2_. These cells were seeded at 50% confluency before 6-Br-Q_0_C_10_ treatment. About 5000 cells/well were seeded in a 24-well plate and treated with various amounts of 6-Br-Q_0_C_10_. Cells were then incubated at 37°C with 5% CO_2_. Cell growth was examined by cell counting stained with Trypan blue. THP-1 cells were cultured in RPMI-1640 medium supplemented with 10% (v/v) heat-inactivated FBS at 37^°^C and 5% CO_2_ (18).

### Cell Attachment Assay

RAW 264.7 and L929 cells were pre-seeded (100,000 cells/well) in 12-well plates (overnight incubation) at 37^°^C and 5% CO_2_. 6-Br-Q_0_C_10_ was then added at 24 hr post pre-seeding, and cell attachment was determined by counting attached cells at 72 hr post treatment. For THP-1 cells, pre-seed 100,000 cells/well in the presence of 320 nM phorbol-12-myristate-13-acetate (PMA) in 12-well plates (overnight incubation) at 37^°^C and 5% CO_2_. 6-Br-Q_0_C_10_ was added at 24 hr post pre-seeding and cell attachment was determined by counting attached cells at 72 hr post treatment.

### MTT Cell Growth Inhibition Assay

HEK293 and HeLa cells were cultured in DMEM supplemented with 10% (v/v) fetal bovine serum and 1% (v/v) penicillin–streptomycin. INS-1 cells were cultured in RPMI 1640 media supplemented with 10% (v/v) FBS and 1% (v/v) penicillin– streptomycin. All the cells were incubated at 37 °C in 5% CO2 atmosphere in 10-cm plates. When cells reached about ∼80 % confluency, they were harvested and cultured on 96-well plates with 5 X 10^3^ cells seeded per well. After 24 hr pre-seeding, the culture medium was replaced by the same mediums containing various concentrations (0, 4, 8, 16, 32 and 64 µM) of 6-Br-Q_0_C_10_. Three replicates were made for each condition. After 24 hr treatment with 6-Br-Q_0_C_10_, the MTT assays were performed using the MTT Assay Kit according to the manufacturer’s manual. Ten μl of the Kit reagent was added into each well, and optical absorption (OD) at 450 nm was measured using a multifunction microplate reader (Infinite M200 Pro, Tecan) after incubation for 2 hr at 37 °C. The cell viability is expressed in the percentage of OD of treated cells verse that of the untreated cells.

### Bisbenzimide-based nuclear fluorescent stain and cell imaging

HEK293 and HeLa cells were prepared and treated with 6-Br-Q_0_C_10_ as described in the section “MTT Cell Growth Inhibition Assay” above. After 24- and 48-hr 6-Br-Q_0_C_10_ treatment, the cells in the 96-well plates were used for bisbenzimide-based nuclear fluorescent stain, and cell imaging. Treated cells were first fixed with 2% paraformaldehyde solution for 20 min at room temperature, and then bisbenzimide (0.5 mg/mL) was added to each well to a final concentration of 1.0 µg/mL. The cells were then incubated at room temperature for 15 min before the cell imaging analysis. Cell images were taken with a Zeiss AXIO Observer. Z1 Fluorescence Motorized Microscope w/ Definite Focus.2 Pred 7. Zen 3.1 (Zen Pro) software was used.

## Results

We first tested the effect of 6-Br-Q_0_C_10_ on the growth of three mammalian cell lines lines (rM-1, ARPE-19 and Hep G2) in cell culture. When these cell lines were cultured for 48 hr in their corresponding media in the presence of various concentrations of 6-Br-Q_0_C_10_ (0, 0.5, 2.5. 5 and 10 µM), the number of live cells decreased as the concentration of 6-Br-Q_0_C_10_ increased (**Table 2**). The effects of 6-Br-Q_0_C_10_ on cell growth of rCM-1 were very similar to that of ARPE-19. Their growths were 6-Br-Q_0_C_10_ concentration dependent. The growth rate of rCM-1 and ARPE-19 cell lines decreased to 20-30% when concentration of 6-Br-Q_0_C_10_ reached 10 µM in cell culture. Under the same conditions, no significant effect on cell growth was observed when normal coenzyme Q or Q_0_C_10_ were used instead of 6-Br-Q_0_C_10_ (Data not shown).

**Table 2.** Effect of 6-Br-Q_0_C_10_ on the growth of rMC-1, ARPE19 and Hep-G2.

| Br-Q<sub>0</sub>C<sub>10</sub> | 0 μM | 0.5 μM | 1 μM | 2.5 μM | 5 μM | 10 μM |
| --- | --- | --- | --- | --- | --- | --- |
| rMC-1 | 100% (1.2x10<sup>6</sup>) | 88.2 | 80.1 | 68.1 | 36.2 | 27.9 |
| ARPE19 | 100% (3.1x10<sup>5</sup>) | 85.4 | 80.9 | 60.0 | 41.9 | 26.6 |
| Hep G2 | 100% (1.3x10<sup>6</sup>) | 103.1 | 107.8 | 104.2 | 98.1 | 93.3 |

The effect of 6-Br-Q_0_C_10_ on Hep-G2 was minimal, even at a high (10 µM) concentration of 6-Br-Q_0_C_10_. One possible explanation is that Hep-G2 cells might have high concentration of native coenzyme Q, a higher concentration of 6-Br-Q_0_C_10_ would be needed in order to compete against coenzyme Q to have higher inhibition.

We further tested the effect of 6-Br-Q_0_C_10_ on the growth of three other cell lines in cell culture, including mouse fibroblast (L929), mouse macrophages (RAW 264.7) and human monocytes (THP-1). All cells were cultured under the same conditions with their corresponding media (as shown in **Table 1**) (18-20) in the presence of 6-Br**-**Q_0_C_10_. The concentrations of 6-Br-Q_0_C_10_ used were: 0, 5, 10, 20, 30 and 50 µM, as shown in Figure 3. As expected, 6-Br-Q_0_C_10_ inhibited cell growth in a concentration-dependent manner. A 50% inhibition of cell growth (Figure 3A) was observed when the concentration of 6-Br-Q_0_C_10_ reached 50 µM. Under the same condition, a 70% inhibition (Figure 3B) was observed for the attached cells.

**Figure 3.**
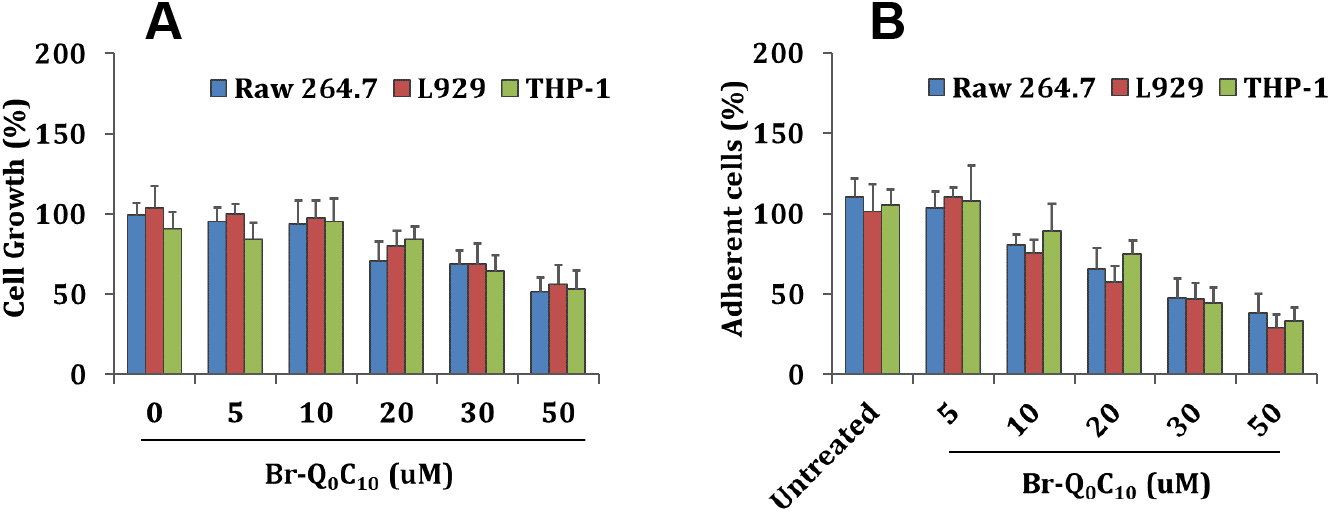
The effect of 6-Br-Q_0_C_10_ on cell growth of L-929, Raw 264.7 and THP1 cells. (A) The inhibition of cell growth with increased concentrations of 6-Br-Q_0_C_10_. (B) The inhibition of attached cells with increased concentrations of 6-Br-Q_0_C_10_.

Figure 4 shows the effects of 6-Br-Q_0_C_10_ on another set of cell lines (THP-1, HEK-293, HELA and ADSC), using three detecting methods—cell counting (Fig 4A), MTT (Fig 4B) and cell imaging (Fig 4C). As described in the previous sections, cell lines were cultured in their corresponding media (**Table 1**), in present of various concentration of 6-Br-Q_0_C_10_ (0, 2, 4, 8,16, 32 and 64 µM) for 48 and 72 hrs. After 48 hours treatment, inhibitions of 69, 66 and 82% were observed for HEK-293, HELA and ADSC (Figure 4A), respectively, at a concentration of 64 μM of 6-Br-Q_0_C_10_. Similar results were obtained in the MTT assay (Figure 4B). As expected, when the incubation prolonged (72 hrs), more inhibitions were observed. A respective inhibition of 95%, 97%, and 90% was observed in HEK-293, HELA, and ADSC cell lines. The effect of 6-Br-Q_0_C_10_ on INS-1 cell line, however, was much less.

**Figure 4.**
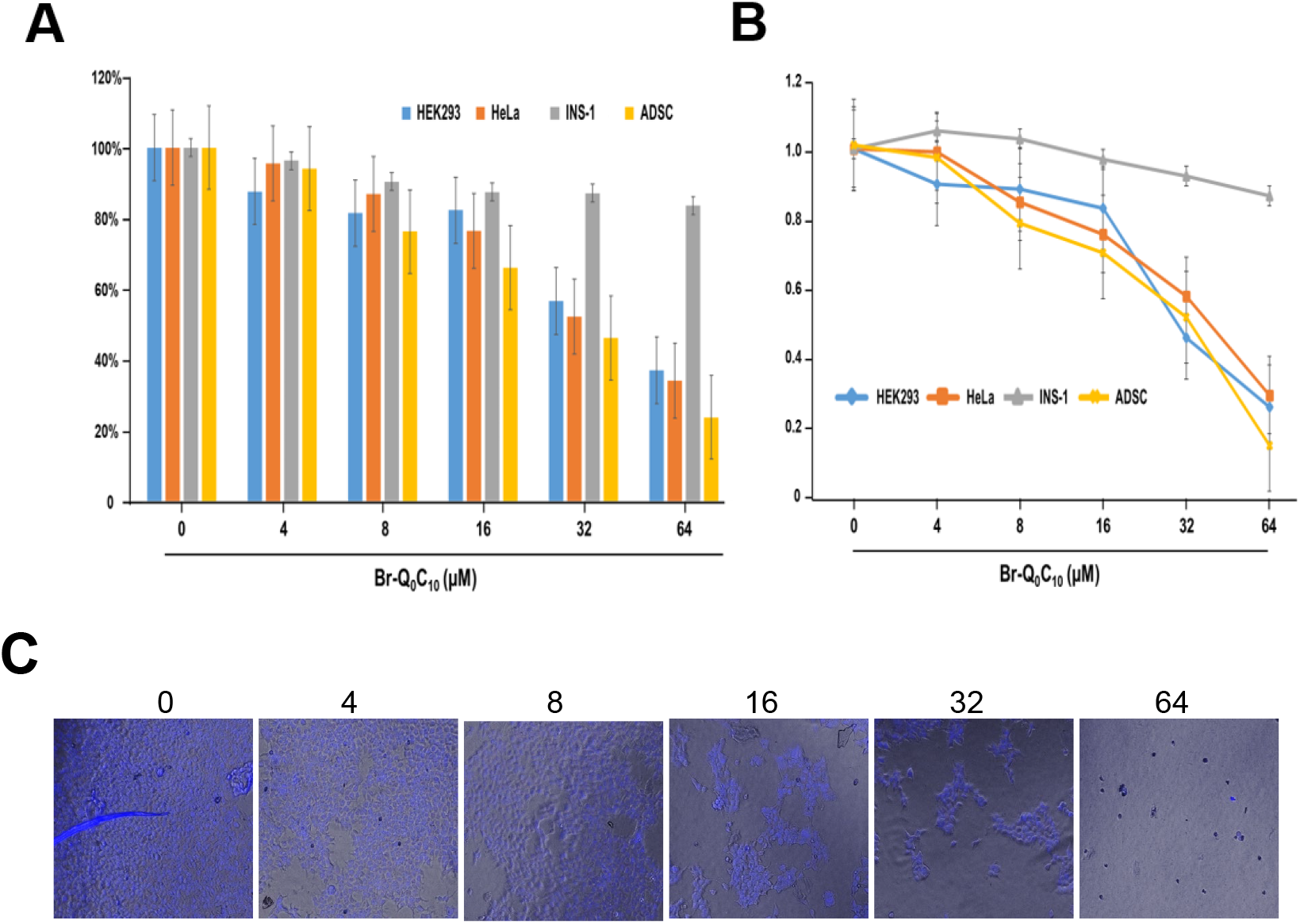
The effect of 6-Br-Q_0_C_10_ on cell growth of HEK293, HeLa, INS-1 and ADSC. (A) The inhibition of cell growth with increased concentrations of 6-Br-Q_0_C_10_ measured by counting viable and nonviable cells using Trypan Blue and a haemocytometer. (B) Cell Growth Inhibition determined by MTT cytotoxicity assay. (C) Representative images of HEK293 cells after 48 hours treatment with increased concentrations of 6-Br-Q_0_C_10._ Cells were stained with bisbenziminde.

## Discussions

As described in the Materials and Methods section, multiple cell-lines, analytical methods, and independent laboratories were involved in this investigation in order to assure the observations made are general occurrence and not due to any specific detecting condition or a specific cell-line. It should be noted that the decrease in cell growth does not appear to result from toxicity a brominated metabolite of 6-Br-Q_0_C_10_ because a similar decrease in cell growth was also observed when 6-Cl-Q_0_C_10_ was used. This finding suggests that the growth-inhibitory effect is likely attributable to modulation of mitochondrial bioenergetics rather than compound-specific cytotoxicity.

Although 6-Br-Q_0_C_10_ inhibits cell growth of all the cell lines tested, its effectiveness does vary slightly among different cell lines. This variability could be due to the different concentration of Coenzyme Q in the different cell lines. The effect of 6-Br-Q_0_C_10_ on cell growth might be enhanced if it is used together with the cholesterol lowing drugs such as Crestor or Lipitor, as they are also known to decrease the biosynthesis of coemzyme O_10_. It is conceivable that fast-growing cells (such as cancer cells) will be more affected by 6-Br-Q_0_C_10_ than the slower growing ones. Restricting the energy supply to the cancer cells might slow down their growth and thus prolong patient’s life. The experiments on the effect of 6-Br- or 6-Cl-Q_0_C_10_ on the weight gain of rats are currently in progress.

Ongoing studies are evaluating the effects of 6-Br-Q_0_C_10_ and 6-Cl-Q_0_C_10_ on weight gain in rat models. In collaboration with Professor Su-lie Chang at Sedan Hall University, preliminary observations indicate a significant reduction in weight gain in rats fed a high-fat diet (personal communication). Further investigation is warranted to validate these findings and to assess long-term metabolic outcomes.

## Acknowledgement

This work was supported by grants GM30721 from the National Institutes of Health, and Oklahoma State University.

## References

1. Biner O, Schick T, Ganguin AA, von Ballmoos C. Towards a Synthetic Mitochondrion. Chimia (Aarau). 2018;72(5):291–6. doi: 10.2533/chimia.2018.291. PubMed PMID: 29789065.

2. Alcázar-Fabra M, Navas P, Brea-Calvo G. Coenzyme Q biosynthesis and its role in the respiratory chain structure. Biochim Biophys Acta. 2016;1857(8):1073–8. Epub 20160310. doi: 10.1016/j.bbabio.2016.03.010. PubMed PMID: 26970214.

3. Trumpower BL, Gennis RB. Energy transduction by cytochrome complexes in mitochondrial and bacterial respiration: the enzymology of coupling electron transfer reactions to transmembrane proton translocation. Annu Rev Biochem. 1994;63:675–716. doi: 10.1146/annurev.bi.63.070194.003331. PubMed PMID: 7979252.

4. Lerner KL, Nemeh K, Longe J. The Gale Encyclopedia of Science, 6th edition (K. Lee Lerner, Contributing Advisor): Gale; 2021.

5. Sazanov LA, Hinchliffe P. Structure of the hydrophilic domain of respiratory complex I from Thermus thermophilus. Science. 2006;311(5766):1430–6. Epub 20060209. doi: 10.1126/science.1123809. PubMed PMID: 16469879.

6. Hirst J. Mitochondrial complex I. Annu Rev Biochem. 2013;82:551–75. Epub 20130318. doi: 10.1146/annurev-biochem-070511-103700. PubMed PMID: 23527692.

7. Sun F, Huo X, Zhai Y, Wang A, Xu J, Su D, et al. Crystal structure of mitochondrial respiratory membrane protein complex II. Cell. 2005;121(7):1043–57. doi: 10.1016/j.cell.2005.05.025. PubMed PMID: 15989954.

8. Cecchini G. Function and structure of complex II of the respiratory chain. Annu Rev Biochem. 2003;72:77–109. doi: 10.1146/annurev.biochem.72.121801.161700. PubMed PMID: 14527321.

9. Xia D, Yu CA, Kim H, Xia JZ, Kachurin AM, Zhang L, et al. Crystal structure of the cytochrome bc1 complex from bovine heart mitochondria. Science. 1997;277(5322):60–6. doi: 10.1126/science.277.5322.60. PubMed PMID: 9204897; PubMed Central PMCID: PMC12235523.

10. Iwata S, Lee JW, Okada K, Lee JK, Iwata M, Rasmussen B, et al. Complete structure of the 11-subunit bovine mitochondrial cytochrome bc1 complex. Science. 1998;281(5373):64–71. doi: 10.1126/science.281.5373.64. PubMed PMID: 9651245.

11. Senior AE, Nadanaciva S, Weber J. The molecular mechanism of ATP synthesis by F1F0-ATP synthase. Biochim Biophys Acta. 2002;1553(3):188–211. doi: 10.1016/s0005-2728(02)00185-8. PubMed PMID: 11997128.

12. Shinzawa-Itoh K, Sugimura T, Misaki T, Tadehara Y, Yamamoto S, Hanada M, et al. Monomeric structure of an active form of bovine cytochrome c oxidase. Proc Natl Acad Sci U S A. 2019;116(40):19945–51. Epub 20190918. doi: 10.1073/pnas.1907183116. PubMed PMID: 31533957; PubMed Central PMCID: PMC6778200.

13. Crane F, Barr R. Chemical structure and properties of coenzyme Q and related compounds. Coenzyme Q biochemistry, bioenergetics and clinical applications of ubiquinone John Wiley & Sons, Chichester, United Kingdom. 1985:1–37.

14. Yu CA, Yu L. Syntheses of biologically active ubiquinone derivatives. Biochemistry. 1982;21(17):4096–101. doi: 10.1021/bi00260a028. PubMed PMID: 6289870.

15. Shunk CH, Linn BO, Wong EL, Wittreich PE, Robinson FM, Folkers K. Coenzyme Q. II. SYNTHESIS OF 6-FARNESYL- AND 6-PHYTYL-DERIVATIVES OF 2,3-DIMETHOXY-5-METHYLBENZOQUINONE AND RELATED ANALOGS. Journal of the American Chemical Society. 1958;80(17):4753-. doi: 10.1021/ja01550a097.

16. Yu CA, Li, X.L., Gu, L.Q., Yu L. Interaction of 6-Bromo- and 6-Chloro-Uniquinone Deivative with the Mitochondrial Electron Transport System. Journal of Chemistry. 2024;10(1):7. doi: 10.17352/ojc.000036.

17. Haslam DW, James WP. Obesity. Lancet. 2005;366(9492):1197–209. doi: 10.1016/s0140-6736(05)67483-1. PubMed PMID: 16198769.

18. Guthrie CM, Meeker AC, Self AE, Ramos-Leyva A, Clark OL, Kotey SK, et al. Microvesicles Derived from Human Bronchial Epithelial Cells Regulate Macrophage Activation During Mycobacterium abscessus Infection. J Proteome Res. 2025;24(5):2291–301. Epub 20250328. doi: 10.1021/acs.jproteome.4c00827. PubMed PMID: 40153482; PubMed Central PMCID: PMC12053935.

19. Vermeire CA, Tan X, Ramos-Leyva A, Wood A, Kotey SK, Hartson SD, et al. Characterization of Exosomes Released from Mycobacterium abscessus-Infected Macrophages. Proteomics. 2025:e202400181. Epub 20240916. doi: 10.1002/pmic.202400181. PubMed PMID: 39279549.

20. Kotey SK, Tan X, Fleming O, Kasiraju RR, Dagnell AL, Van Pelt KN, et al. Intracellular iron accumulation facilitates mycobacterial infection in old mouse macrophages. Geroscience. 2023. Epub 20231230. doi: 10.1007/s11357-023-01048-1. PubMed PMID: 38159133.

21. Emmerich SD, Fryar CD, Stierman B, Ogden CL. Obesity and Severe Obesity Prevalence in Adults: United States, August 2021-August 2023. NCHS Data Brief. 2024(508). doi: 10.15620/cdc/159281. PubMed PMID: 39808758; PubMed Central PMCID: PMC11744423.

